# Optimal antimicrobial dosing combinations when drug-resistance mutation rates differ

**DOI:** 10.1101/2024.05.04.592498

**Authors:** Oscar Delaney, Andrew D. Letten, Jan Engelstädter

## Abstract

Given the ongoing antimicrobial resistance crisis, it is imperative to develop dosing regimens optimised to avoid the evolution of resistance. The rate at which bacteria acquire resistance-conferring mutations to different antimicrobial drugs spans multiple orders of magnitude. By using a mathematical model and computer simulations, we show that knowledge of relative mutation rates can meaningfully inform the optimal combination of two drugs in a treatment regimen. We demonstrate that under plausible assumptions there is a linear relationship in log-log space between the drug *A*:drug *B* dose ratio that maximises the chance of treatment success and the ratio of their mutation rates. This power law relationship holds for bacteriostatic and bactericidal drugs. If borne out empirically, these findings suggest there might be significant room to further optimise antimicrobial dosing strategies.

## 1 Introduction

One of the key goals of designing antimicrobial treatment regimens must be to minimise the probability that resistance develops, alongside striving to rapidly clear the patient’s infection and avoid excessive toxicity. One valuable approach is to use multiple drugs, either in combination [1, 2] or sequentially [3, 4, 5, 6] such that even if a mutation conferring resistance to a single drug occurs, the mutant is still impacted by the other drug(s). Using multiple drugs, rather than just a larger dose of a single drug, may also reduce toxic side effects in the patient, especially if the drugs interact synergistically and hence allow for smaller concentrations to be efficacious [7]. Combination therapy is supported by a significant body of empirical literature (reviewed in [8] and [9]), with positive results for example in laboratory evolution settings [1] and tuberculosis treatment [10]. A meta-analysis involving 4514 patients from 53 studies of multidrug-resistant gram-negative bacterial infections found an average reduction in mortality of 17% with combination compared to monotherapy [11].

Many mathematical and computational models have been created to better understand and predict the evolution of resistance (reviewed in [12, 13]). In most models, a key parameter is the mutation rate: the probability that a cell division in a susceptible bacterium will give rise to a cell resistant to the drug in question. The higher the mutation rate, the more likely resistance is to develop (setting aside resistance arising from horizontal gene transfer). Because this relationship is trivial, in most mathematical models the mutation rate is fixed and then ignored, and other putatively more interesting phenomena are explored [14, 15, 16, 17, 18]. Here, we show that in combination therapy, the relative mutation rates for each drug can be an important factor in choosing the optimal quantity of each drug to apply. This differs notably from the conventional wisdom that it is often best to use equal doses of two drugs (e.g. [19]).

An important consideration in combination therapy is whether to use bacteriostatic drugs (i.e. drugs that inhibit growth), bacteriocidal drugs (i.e. drugs that kill bacteria), or both. Theoretical work has shown that when only one drug is present at a time, bacteriostatic drugs are usually more effective in minimising resistance evolution [20]. When two drugs are used in combination, theory suggests that pairing a bacteriostatic drug and a bactericidal drug is especially effective at both clearing the infection and reducing the probability of resistance evolving [19]. Moreover, the density-dependence and resource limitations of the bacterial population impact the relative efficacy of different drug modes of action [21].

The rate at which resistance mutations to a given antimicrobial drug occur may depend on the bacterium that is targeted, the current resource availability or other environmental conditions. Even within one host species and constant conditions, resistance mutation rates can vary greatly by drug [22]. This is unsurprising, as different mechanisms of action may be more or less difficult for the bacteria to surmount or circumvent when sampling from the space of possible mutations. The distribution of fitness effects of possible resistance mutants can also vary greatly by type of drug used [23]. In *Mycobacterium tuberculosis*, the infectious agent responsible for the most deaths per year worldwide [24], there is an approximately 400-fold difference in the mutation rate between two of the most commonly used first-line drugs, rifampicin and ethambutol [25]. That said, it is difficult to accurately compare estimated mutation rates for different drugs. This is because the mutation rate may depend on the drug concentration used (the higher the concentration, the fewer resistance mutations may be possible), and also because the stress response to some drugs may elevate the mutation rate itself [26]. To our knowledge, there is no centralised database of mutation rate estimates, but some example values from the literature for various drugs and species are provided in Table 1. Resistance mutation rates being orders of magnitude apart could reasonably be expected to prove important when choosing optimal dosing strategies. Intuitively, all else being equal, it is better to use drugs for which resistance mutations arise at a lower rate to minimise the probability of resistance developing.

**Table 1:**
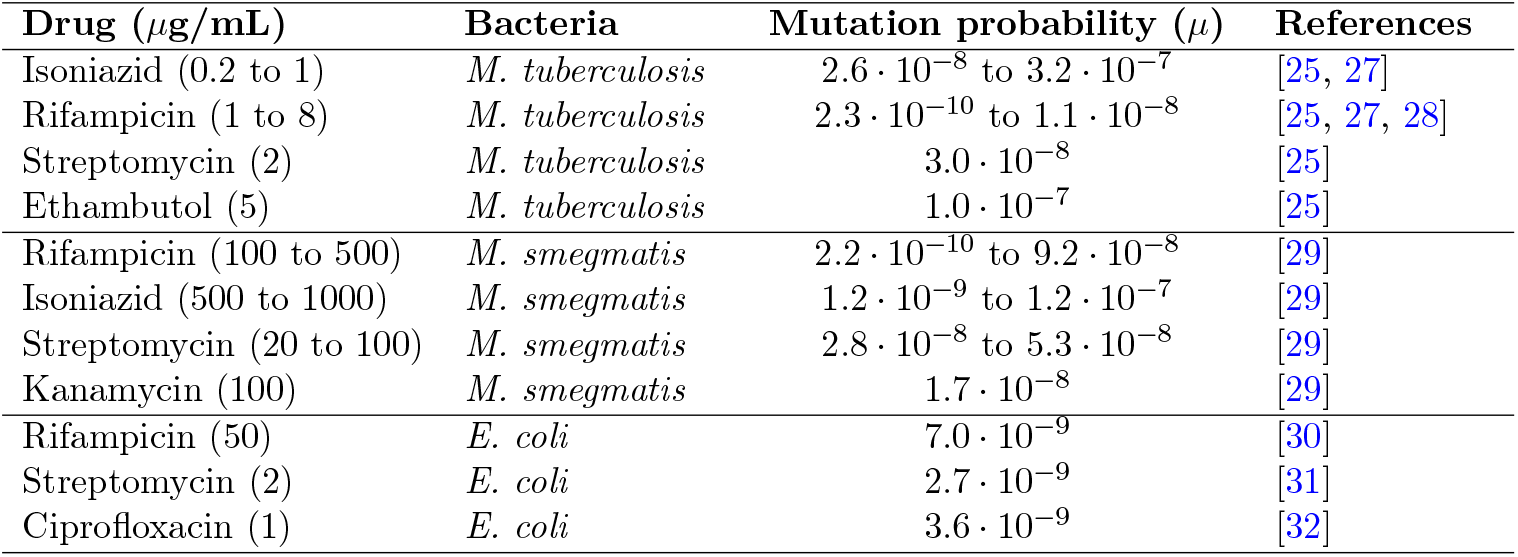
Genome-wide probability of a resistance mutation per replication for various antibiotics in *Mycobacterium tuberculosis, Mycobacterium smegmatis*, and *Escherichia coli*.

We formalise and interrogate this intuition under a variety of plausible assumptions, and develop theoretical predictions for how different resistance mutation rates should alter optimal dosing strategies. We find that a quadrupling of the ratio of mutation rates leads to a doubling in the optimal drug dosing concentration ratios favouring the less evolvable drug. This power law relationship is qualitatively robust to relaxing various simplifying assumptions.

## 2 Methods

We modelled a simple scenario where there is one species of bacteria and two arbitrary drugs, *A* and *B*, administered in combination at concentrations *C*_*A*_ and *C*_*B*_ that are constant over time (we later relax this assumption). After *t* hours the sizes of the susceptible, *A*-resistant, and *B*-resistant populations respectively are *S*(*t*), *M*_*A*_(*t*) and *M*_*B*_(*t*).

To model drug mode of action and pharmacodynamics, we normalised the effective drug concentration (*E*_*i*_) for bacterial strain *i* ∈ {*S, M*_*A*_, *M*_*B*_} (that is, susceptible, *A*-resistant mutants, and *B*-resistant mutants respectively) and drug *j* ∈ {*A, B*} onto the [0, 1) interval using the sigmoid E_max_ model [33] (closely related to the more common Hill equation [34]). Here, *z*_*i,j*_ is the drug concentration at which the half-maximal effect of drug *j* is achieved in strain *i* (denoted EC_50_ in [35]) and *β* is the shape parameter which determines the steepness of the function around *z* [36]:

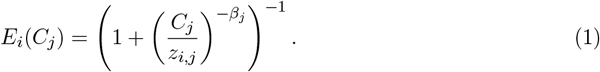

Here, complete resistance (*E*_*i*_(*C*_*j*_) = 0 for any *C*_*j*_) would arise with 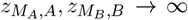. We consider drugs that are either bacteriostatic (denoted *ϕ*_*j*_ = 1) and only affect the cell division rate, or bactericidal (denoted *ϕ*_*j*_ = 0) and only affect the cell death rate, leaving out intermediate cases. We ignored the effect of intra-specific competition on growth, such that the replication rate of strain *i* is a constant *r*_*i*_ in the absence of drugs (this assumption is later relaxed in the simulations, but is necessary for analytical progress). While in most cases bacterial growth is resource-limited such that our analytical model would be unrealistic, in some cases e.g. if antibiotic treatment is started early by the human host before pathogenic bacteria have reached the resource limits of their niche, our model could be more realistic. Likewise, *δ*_*i*_ is a constant intrinsic death rate term, representing constant negative pressures from competition with other (non-modelled) bacterial species [37], and the host’s immune response. Combining, the drug-dependent replication, death, and net growth rates are

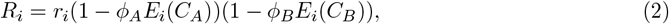

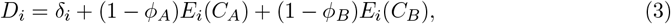

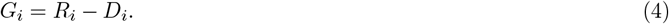

Our model is based on ‘Bliss independence’ (introduced in [38]) which assumes that the two drugs have distinct, independent, cellular targets and modes of action [39]. In the case of bactericidal drugs, the null model of no synergistic or antagonistic drug interaction is given by the total mortality rate from the drug combination equalling the sum of the mortality rates that would ensue with each drug used in isolation. However, for bacteriostatic drugs, a null interaction means that each drug reduces the replication rate by the same factor in combination as when used in isolation. For a cell to die, it is sufficient for *either* drug to cause its death (akin to a logical OR gate), so these terms are added, whereas for a cell to divide *both* pathways impacted by the drugs must remain functional (akin to a logical AND gate), so the terms are multiplied. While other modeling choices were possible, this approach is reasonable as successful cell division requires each necessary pathway to be functioning, and thus the probabilities of each pathway being functional should be multiplied.

We denote the probability of a cell division event leading to a *j*-resistant daughter cell as *µ*_*j*_, and ignore back-mutations and the (initially negligible) chance of double-mutations. We initially assume that the mutation rate is independent of the drug concentration used, though this assumption is later relaxed. Thus, a deterministic version of our model can be represented as the following system of ordinary differential equations (ODEs):

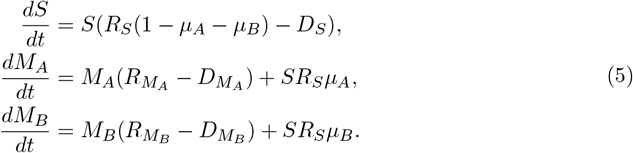

Along with the initial conditions, where only susceptible cells are present (*S*(0) = *S*_0_, *M*_*A*_(0) = 0, *M*_*B*_(0) = 0), this fully defines the mathematical model. To more realistically model the uncertainty inherent in growth and mutation, we employed a stochastic version of this model. Specifically, for the computational implementation, we used the Stochastic Simulation Algorithm (also known as the Gillespie algorithm [40]) to evolve the system over time, with birth and death events given in Table 2. All simulations were performed using R v4.3.0 [41]. To make evolving this stochastic system less computationally intensive, we used tau-leaping to perform many transitions in one step with the *adaptivetau* package [42, 43]. We used the *future* package to parallelise simulation runs [44]. To store, analyse, and visualise the simulation data we used the *tidyverse* set of packages [45, 46].

**Table 2:** Transition events and rates for the Gillespie algorithm.

| Event | Transition | Rate |
| --- | --- | --- |
| S birth | $S \rightarrow S + 1$ | $SR_S(1 - \mu_A - \mu_B)$ |
| $M_A$ birth | $M_A \rightarrow M_A + 1$ | $SR_S\mu_A + M_AR_{M_A}$ |
| $M_B$ birth | $M_B \rightarrow M_B + 1$ | $SR_S\mu_B + M_BR_{M_B}$ |
| S death | $S \rightarrow S - 1$ | $SD_S$ |
| $M_A$ death | $M_A \rightarrow M_A - 1$ | $M_AD_{M_A}$ |
| $M_B$ death | $M_B \rightarrow M_B - 1$ | $M_BD_{M_B}$ |

The value we seek to maximise is the probability that the susceptible population is driven to extinction without resistance becoming established. This can be operationalised as the probability that *S*(*t*) +*M*_*A*_(*t*) +*M*_*B*_(*t*) = 0 for any time *t*. Trivially, arbitrarily large drug concentrations are optimal for this goal. However, toxic side effects for the host mean that drug concentrations must be restricted. We use a simple toxicity model with some fixed maximum allowable toxicity *c*, and both drugs contribute equally and linearly to this maximum, that is *C*_*A*_ + *C*_*B*_ ≤ *c*. To maximise the combined efficacy of the drugs, the highest allowable concentrations are used (*C*_*A*_ + *C*_*B*_ = *c*).

The drugs are assumed to be equally effective, and their concentrations are scaled to be in units standardised to the potency of the drug in question, such that *z*_*S,A*_ = *z*_*S,B*_ = 1. We initially assume complete resistance 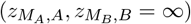 and a default of no cross-resistance or collateral sensitivity such that 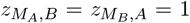. We use a default value of the maximum replication rate of *r*_*i*_ = 1 h^−1^, and of the (dimensionless) Hill coefficient of *β* = 1, using convenient round numbers that are realistic for some bacteria and drugs [47]. Denoting the total chance of a mutation conferring resistance as *µ* = *µ*_*A*_ + *µ*_*B*_, we use *µ* = 10^−9^ per replication which is in the range of common values in Table 1. We use a starting population size of *S*_0_ = 10^9^ cells and an intrinsic death rate of 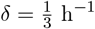, which are plausible biological values [48]. Common values of the maximum drug-induced death rate of bactericidal drugs are anywhere from approximately 1 h^−1^ to 10 h^−1^ [47]. However, the theoretical maximum efficacy of a bacteriostatic drug is 1, that is preventing 100% of replications. To avoid skewing the model towards bactericidal drugs, we use a default value of 1 for the maximum drug-induced death rate too, which is at the lower end of common values. We chose all these parameter values first and foremost to ensure resistance evolves a middling fraction of the time - resistance occuring almost never or always would not yield biologically interesting phenomena - while using roughly plausible round values, rather than focusing on precise estimates from the empirical literature.

Holding all other parameters constant, we seek a mapping (*µ*_*A*_, *µ*_*B*_) → (*C*_*A*_, *C*_*B*_) that maximises the probability of eventual extinction, *P*_*E*_. We call a *strategy* a choice of what drug concentrations *C*_*A*_ ∈ [0, *c*] and *C*_*B*_ = *c* − *C*_*A*_ to apply. Drugs for which resistance mutations arise at a lower rate are preferred, however the diminishing marginal returns to increasing drug concentrations defined by the *E*_*i*_(*C*_*j*_) function mean that it is not necessarily optimal to use only the drug with a lower mutation rate.

## 3 Results

### 3.1 Analytical solution

To find the probability *P*_*D*_ that a newly arisen resistant mutant cell leaves no descendants in the distant future, we can use the law of total probability, noting that a mutant is either resistant to *A* or *B*. Denoting the probability a strain-*i* mutant cell leaves no descendants in the distant future as *P*_*D*|*i*_, we get

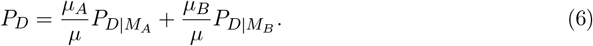

Due to the stochastic nature of the model, even a mutant lineage with a positive growth rate may become extinct, and thus *P*_*D*|*i*_ is not necessarily 0. We can again use the law of total probability, noting that the cell must either die before dividing or divide before dying, and that the probability of each occurring first is proportional to the rate of that stochastic process. If the cell successfully divides once, each of the two daughter cells will also have a *P*_*D*|*i*_ chance of leaving no descendants, as they are functionally identical and independent. This gives

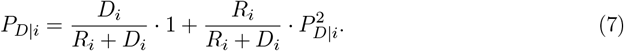

This is a special case of the well-characterised Gambler’s Ruin problem, where a bettor repeatedly wagers to either gain or lose a single dollar, until either they have lost everything, or reached some target value. In our case, extinction occurs if and only if every lineage initiated by a single mutant cell eventually dies out, and the solution known since Fermat [49] is that

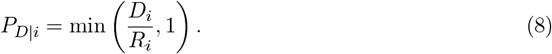

Now we can approximate the number of cell divisions (𝒩) in the susceptible population as a deterministic process, as it begins with a very large number of cells so the stochasticity of individual cell divisions becomes negligible. Given that *µ* ≪ 1 (see e.g. Table 1), we can ignore losses from mutation and thus use the approximation 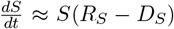 and therefore 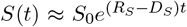. Given that under antibiotic treatment *R*_*S*_ < *D*_*S*_ and hence *G*_*S*_ < 0, this means the susceptible population undergoes exponential decay. We can then estimate the total number of replications 𝒩 as

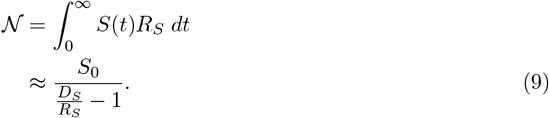

The probability that each cell starts a successful resistant lineage is the product of the probability of a resistance mutation (*µ*) and the probability that a resistant mutant leaves descendants (1 − *P*_*D*_). Noting that the outcome of each new cell is independent, we find the overall extinction probability is

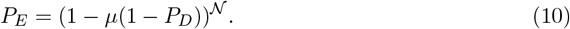

Because *µ* ≪ 1, we can again make the approximation

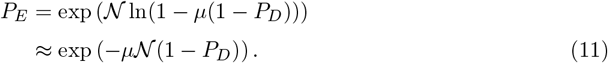

This finding is structurally very similar to the classic result from the evolutionary rescue theory literature that *P*_*E*_ ≈ exp (−*N*_0_ *θ*) where *N*_0_ is the initial population introduced to a novel environment, and *θ* is the rate of rescue for each individual [50]. In our case, 𝒩 replaces *N*_0_ given the relevant quantity is the number of replications, not the inoculum size, and *θ* is replaced by the probability a mutation occurs and survives, *µ*(1 − *P*_*D*_).

Equation 11 will be used for the computational implementation, as complicated expressions are unproblematic for numerical methods. But to make further analytical progress, it is useful to simplify the analysis by considering a small class of possible parameters that make the formulas collapse down to more manageable forms. In particular, the simplifying assumptions are:

- Resistant cells are unaffected by arbitrarily high drug concentrations 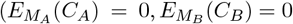 for all drug concentrations *C*_*A*_, *C*_*B*_ ∈ ℝ ^+^).
- The shape parameters of the pharmacodynamic functions are unity (*β*_*A*_ = *β*_*B*_ = 1, equivalent to Michaelis-Menten kinetics).
- The drug-free replication rate and death rate of all strains are the same (that is, there is no cost of resistance: *r*_*i*_ = *r, δ*_*i*_ = *δ*).
- Both drugs are bacteriostatic (*ϕ*_*A*_ = *ϕ*_*B*_ = 1).
- When only drug *A* is applied (*C*_*A*_ = *c, C*_*B*_ = 0), the net growth rate of the susceptible and *B*-resistant strains are both zero 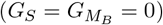, and vice versa for when only drug *B* is applied. (Even with both drugs being bacteriostatic, some replication events can occur, which is why the net growth rate is not negative.) This implies that *r* = *δ*(1 + *c*) and thus that 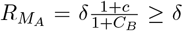 and likewise for *M*_*B*_. Thus, *G*_*S*_ ≤ 0 and 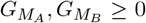. If the toxicity restriction were relaxed, and both drugs are used with a full dose (*C*_*A*_ = *C*_*B*_ = *c*) the population would be eradicated without resistance evolution as neither single-resistant strain could grow.

These simplifying assumptions may in reality often be violated, but they are plausible simplifications. For example, some resistance mutations do confer resistance even to relatively high drug concentrations [51], and costs of resistance can be small [52]. The final assumption is less conceptually important, but makes the computations simpler. While restrictive, these assumptions are still plausible enough to be interesting, and will be relaxed later in the Simulations section. After some algebraic manipulations, substituting these assumptions into equation 11 yields

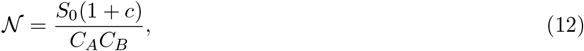

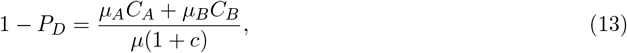

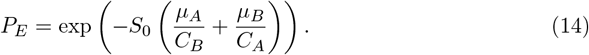

These are pleasingly interpretable equations. 𝒩 is minimised when *C*_*A*_ = *C*_*B*_ given the diminishing marginal efficacy of each drug (Equation 12). Conversely, if only *A* or *B* is used (that is, *C*_*B*_ = 0 or *C*_*A*_ = 0, respectively), 𝒩 is unbounded, as the susceptible population is not killed. The survival probability of a mutant is more sensitive to an increase in concentration of the drug to which resistance mutations arise more frequently (Equation 13). Finally, as *µ*_*A*_ increases relative to *µ*_*B*_ the infection is more likely to be cleared with a higher dose of drug *B* than drug *A*, because *A*-resistant cells are still susceptible to drug *B* (Equation 14).

We maximised *P*_*E*_ by computing its derivative with respect to *C*_*A*_, setting this equal to 0, and solving for the optimal drug concentrations, denoted *Ĉ*_*A*_ and *Ĉ*_*B*_, to find that

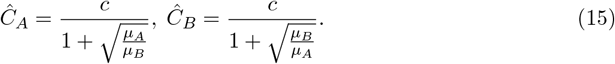

This can also be rewritten in ratio form:

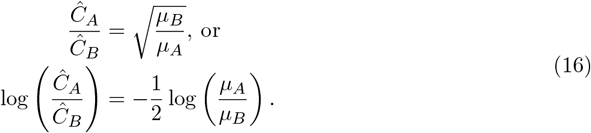

The second version of this equation is useful as it shows that in log-log space there should be a linear relationship between the ratio of the mutation rates and the ratio of the doses. In other words, there is a power law relationship between the ratios of mutation rates and the ratio of optimal drug doses, with an exponent of 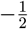. This relationship exhibits the expected behaviour whereby *µ*_*A*_ → 0 entails *Ĉ*_*B*_ → 0 and *µ*_*B*_ → 0 entails *Ĉ*_*A*_ → 0. This means, as mutations become more biased towards conferring resistance to one drug, the optimal combination dosing strategy relies more on the other, less resistance-prone, drug. However, even with a large difference in the mutation rates of the two drugs, the diminishing marginal efficacy of each drug defined by the *E*_*i*_(*C*_*j*_) function means that a nonzero amount of the more resistance-evolution-prone drug should still be used in the drug cocktail.

In the case of both drugs being bactericidal, the intermediate steps are more complicated so are omitted here, but computations in Mathematica v13.1.0.0 [53] show that this same simple relationship in Equation 16 between mutation rates and optimal dosing ratios holds.

### 3.2 Simulations

Here, we corroborate the analytical findings computationally and explore regions of parameter space that appear inaccessible analytically.

When both drugs are bactericidal or both are bacteriostatic, the relationship given in Equation 16 holds (that is, the yellow computational and green theoretical lines coincide in Figure 1A,D). Interestingly, the actual values of *P*_*E*_ differ in the two cases, but the optimal dosing strategy remains the same. When both drugs are bacteriostatic, *P*_*E*_ is lower, as the susceptible population remains large for longer, given there is no drug-induced death (only the intrinsic death rate). A qualitatively similar relationship holds when one drug is bacteriostatic and the other is bactericidal, but the optimal dosing strategy is biased towards the bacteriostatic drug (Figure 1B,C). This effect, where the coupling of mutations to replications favours the growth-inhibiting activity of bacteriostatic drugs, was recently explored in [36].

**Figure 1:**
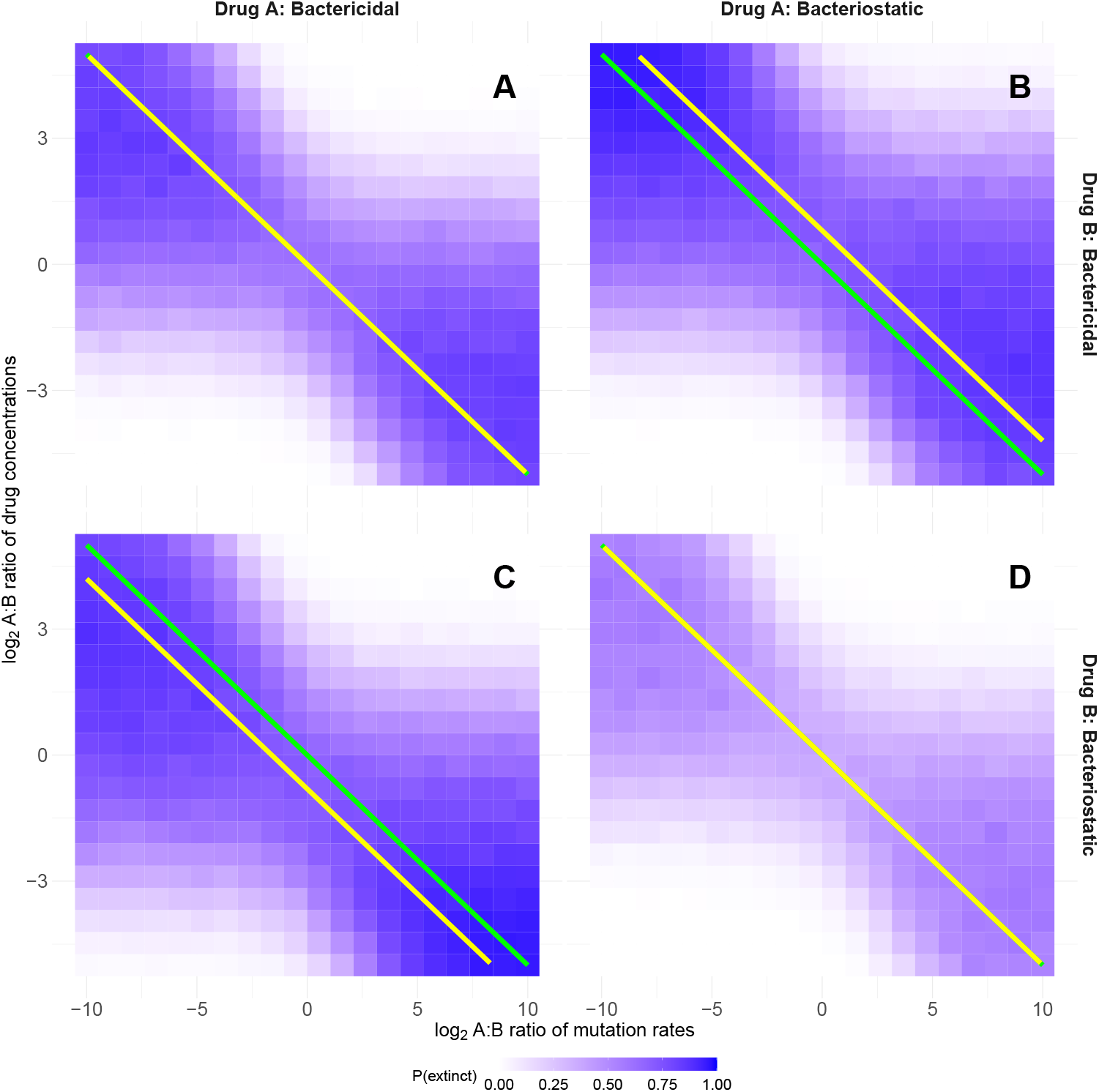
Computational corroboration of basic analytical results. Each grid square shows the probability that an initial population of susceptible bacteria will be driven to extinction by that dosing strategy, averaged over 1000 stochastic simulation runs. The yellow lines show the theoretically optimal dosing strategy for any given ratio of resistance mutation rates, determined by numerically evaluating *P*_*E*_ for many values of *C*_*A*_ and *C*_*B*_ = *c* − *C*_*A*_ using Equation 11, and choosing the minimand and minimum. The green lines are the same in all panels and show the analytical result from Equation 16 for the basic scenario, for comparison. Parameter values are 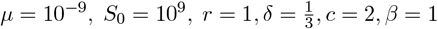 and the drug modes of action vary in each panel. A) *ϕ*_*A*_ = *ϕ*_*B*_ = 0. B) *ϕ*_*A*_ = 1, *ϕ*_*B*_ = 0. C) *ϕ*_*B*_ = 1, *ϕ*_*A*_ = 0. D) *ϕ*_*A*_ = *ϕ*_*B*_ = 1.

Having verified the basic analytical findings, we can begin relaxing various assumptions. We first relax the unrealistic assumption of complete resistance (where 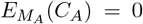 and 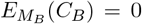 for all drug concentrations, implying 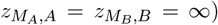. In reality, each resistant strain is still affected by high drug concentrations to some varying extent. To model this, for each simulation run we pick resistance values from a shifted exponential distribution 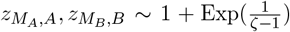 with mean *ζ* > 1. The shift ensures that resistant strains are always more resistant than the susceptible strain 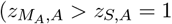 and 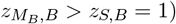. (In reality, some mutations would increase drug susceptibility, but these are not modelled as they would be quickly eliminated from the population.) The exponential shape of the distribution reflects that there are many potential mutations conferring weak resistance and fewer mutations conferring strong resistance [51]. Under this model, some mutants will have low 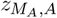 or 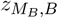 values that entail negative growth rates for therapeutic drug concentrations, making them evolutionary dead ends. In other words, some mutants will sit outside of the mutant-selection window, which effectively reduces the mutation rate by eliminating some fraction of mutations [54].

For a fixed mutation rate ratio and two bacteriostatic drugs, as resistance becomes weaker (*ζ* → 1) the probability of extinction tends towards 1 and the optimal strategy tends towards using equal amounts of both drugs (Figure 2). This is because, given the diminishing marginal efficacy of increased doses of each drug, using equal concentrations minimises the net growth rate and therefore reduces 𝒩, which is most important when the drugs are less effective. Conversely, for strong resistance as *ζ* → ∞, the optimal ratio of drug concentrations converges to the theoretical value given in Eq. 16 (which for the example in Figure 2 computes to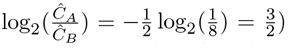. Figure S1 shows that with *ζ* = 5 the results are very similar to those seen in Figure 1. This suggests that our analytical results are still reasonable despite assuming resistance is complete.

**Figure 2:**
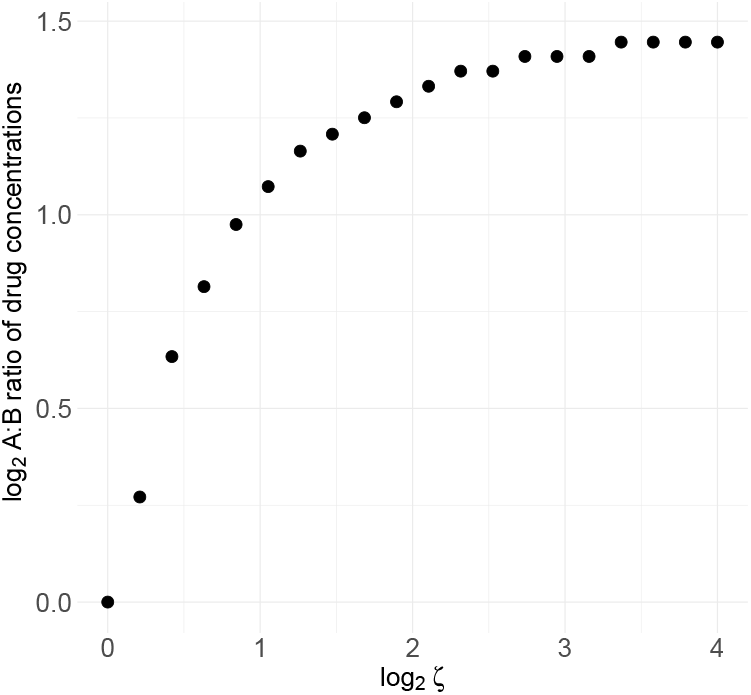
Optimal dosing with partial resistance. Parameters are the same as in Figure 1D except that *µ*_*B*_ = 8*µ*_*A*_ and 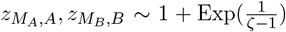 for *ζ* > 1 and 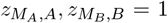 for *ζ* = 1. For each dot defining a *ζ* value, 1000 values of *z* were drawn, evenly spaced from the cdf at the 0.1^*th*^, 0.2^*th*^, …, 99.9^*th*^ percentiles, and the mean probability of extinction was computed over these 1000 using the approximation in Equation 11. This was done for 30 possible ratios of drug doses, and the dosing ratio which yielded the highest *P*_*E*_ value was plotted on the y-axis.

Changing the shape parameter (*β*) noticeably changes the basic result. If *β* > 1 then the pharmacodynamic function has a sigmoidal shape and thus is steeper around the *z*-value where the drug has half its maximal effect. This means that intermediate values of both drugs are less beneficial than a more potent dose of just one drug, especially when both drugs are bacteriostatic. Beyond some threshold *β* value, using just one drug is optimal (Figure 3). At this threshold value the intermediate drug concentration ratio switches from being the global maximum of extinction probability to only a local maximum, so an underlying smooth function leads to a discontinuous result upon taking the maximand. If instead *β* < 1 then the pharmacodynamic function is steep initially near a drug concentration of zero, and then approaches the maximum inhibitory effect slowly. Thus, it is most valuable to use some of both drugs. And again, below some threshold, using equal quantities of both drugs is optimal. Results for different mutation rate ratios and drug types are shown in Figures S2 and S3 for *β* = 3 and *β* = 0.2 respectively.

**Figure 3:**
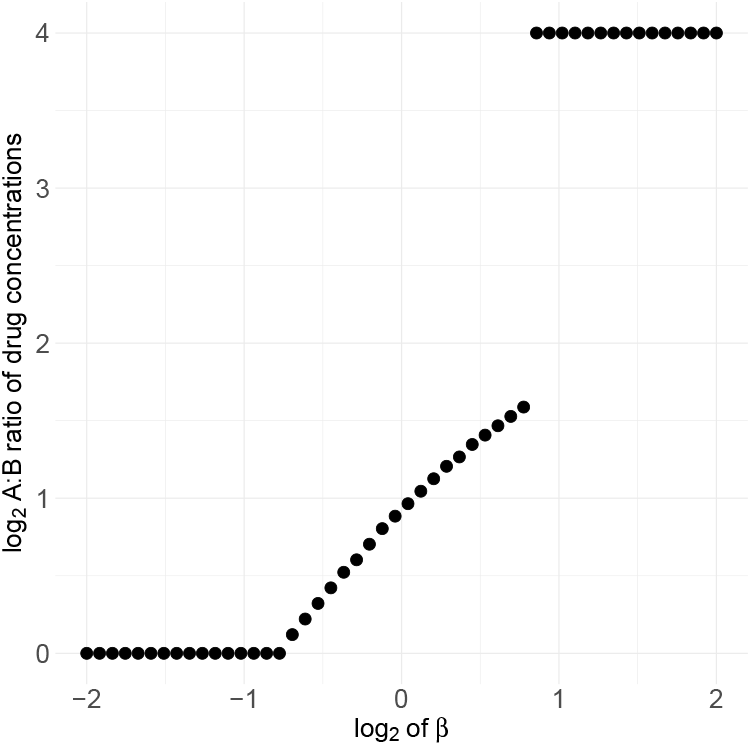
Optimal dosing under shape parameter variation. Parameters are the same as in Figure 1D except that the ratio of mutation rates is fixed at *µ*_*B*_ = 8*µ*_*A*_ and *δ* = 1.1−(1 + *c*^−*β*^)^− 1^ which ensures that regardless of the value of *β* the susceptible population’s growth rate is always at least slightly negative. The maximum drug concentrations ratio tried was 2^4^ = 16, so the fact that dots clump there does not suggest that value is special, instead that arbitrarily large ratios are optimal, but cannot readily be plotted on a finite y-axis.

Thus far cost-free resistance has been assumed, whereas in reality mutations that confer resistance often reduce the maximum replication rate or cause other fitness costs. While some drugs give rise to resistant mutants with unchanged or even increased fitness in the absence of the drug, a meta-analysis suggests that common values of fitness costs are on the order of 10% [52]. For the basic model, in the limit as *C*_*B*_ → 0, *δ* was chosen such that 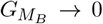, whereas once resistance costs are introduced by setting the replication rate of mutants to be below that of the susceptible strain 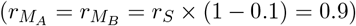 the net growth rate of mutants can become negative. There is a probability of 0 that a mutant with a negative growth rate survives in the long term, and so all negative growth rates are equally good from the perspective of minimising resistance evolution. Thus, here too intermediate dosing strategies are sufficient to ensure *P*_*E*_ ≈ 1 even for very skewed mutation rates (Figure S4). Very skewed drug concentrations also work equally well for very skewed mutation rates - because mutants are less fit in this scenario, there is more flexibility in choosing a drug dosing ratio that results in eradication.

The toxicity-enforced limit of the total drug concentration has so far been fixed at *c* = 2, but this limit is not biologically or theoretically special. We also considered a scenario where the drugs are somewhat less toxic, and a larger maximum dose of *C*_*A*_ + *C*_*B*_ ≤ *c* = 5 can be applied. Maintaining the assumption from before that *G*_*S*_ ≤ 0 and 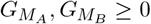 we get that 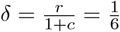 is halved from its earlier value of 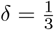. This ensures we still explore an interesting region of parameter space where mutants have a decent chance of arising and surviving. In this case, we observe that the same basic trend holds, while the probability of extinction is higher throughout (Figure S5). The higher overall drug concentrations mean that the optimal strategy skews slightly more heavily towards the bacteriostatic drug (the yellow line is further away from the green line in Figure S5 than in Figure 1) as in absolute terms this still leaves more of the bactericidal drug to clear the infection.

We also extended our basic model to include pharmacokinetics, and found that introducing a drug decay rate of 0.15 h^−1^ left the basic results roughly unchanged (Figure S6). This suggests that ignoring pharmacokinetics in our analytical solution does not undermine its real-world applicability.

Finally, we introduced resource constraints into our basic analytical model [36]. We model the resource-dependent bacterial growth using standard resource-consumption dynamics. Let *R*(*t*) represent the concentration of the resource at time *t* in units standardised such that one cell division uses one unit of resource, and let the resource affinity constant be *K* = 10^8^. The maximum growth rate of strain *i* without drugs is now given by the Monod equation [55] 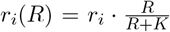 where *r*_*i*_ is the maximum growth rate in resource-unlimited conditions. This creates density-dependent growth that slows as resources become limiting. Let the initial resource level be *ρ* = 10^9^ units, and the constant resource influx rate be *ω* = 10^8^h^−1^. Then resource dynamics follow 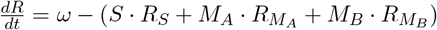 because each replication consumes one unit of resource. We selected parameter values that would allow for a meaningful comparison with our resource-unlimited model. The affinity constant *K* = 10^8^ was chosen to be an order of magnitude below the initial resource concentration (*ρ* = 10^9^), ensuring that growth begins near maximum rates but becomes resource-limited as the population expands. The resource influx rate (*ω* = 10^8^h^−1^) was set to balance consumption when approximately 10^8^ replications occur per hour.

As shown in Figure S7, the fundamental negative relationship between the optimal drug concentration ratio 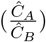 and mutation rate ratio 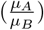 persists under resource-limited conditions, consistent with our theoretical prediction. Though we do not show the theoretical yellow curve for this scenario (as it was derived for resource-unlimited conditions), the computational results still follow approximately the same slope in log-log space. With resource limitation, the susceptible population reaches carrying capacity faster, reducing the total number of replications 𝒩 before antibiotics take effect. This results in fewer mutation events and higher probabilities of extinction across all scenarios compared to the resource-unlimited case. If we set resource constraints to be very limiting, the choice of drugs would matter little, while conversely if resource constraints were very light (e.g. *ρ* = 10^10^) then the results diverge little from the original resource-unconstrained model.

## 4. Discussion

Antimicrobial combination therapy is justified partly on the basis that it reduces the probability of infectious pathogens evolving resistance [1, 2]. To date, however, the design of optimal dosing regimens in combination therapy has given little consideration to drug-specific variation in pathogen resistance mutation rates. Here we have shown that as two drugs have increasingly different mutation rates, the optimal dosing strategy entails using an increasingly large fraction of the drug with a lower resistance mutation rate, according to a simple power law relationship. This is an intuitive result, as drugs that have a higher resistance mutation rate are riskier to use. This result is relatively robust to changing the drugs’ modes of action. Across various alterations to the basic scenario — such as changes to the pharmacodynamic shape parameter *β*, making resistance costly or incomplete, increasing the death rate, or adding pharmacokinetics — the relationship between a skewed mutation rate and skewed optimal dosing strategy persists.

Antibiotic resistance is particularly concerning in tuberculosis, where as of 2022, 12% of all cases worldwide involved multidrug-resistant (MDR) strains of *M. tuberculosis* [56]. A key component of an MDR containment strategy is to minimise the incidence of already resistant strains acquiring resistance to another drug which was previously efficacious. Clinical data from Georgia indicates that for MDR patients being treated with second-line antibiotics, 9% acquire resistance to ofloxacin during treatment, and 10% to kanamycin [57]. To our knowledge, there is no empirical data linking the probability of tuberculosis patients acquiring resistance to a particular drug with the rate at which resistance mutations to that drug arise in the laboratory, however there is a strong prima facie reason to expect such a connection. Mutation rate differences among strains of *M. tuberculosis* have been investigated, and indeed a strain with more frequent mutations in the laboratory had elevated levels of MDR in clinical infections [58].

Our choices of functional forms for the drug-dependent mortality and replication rates in Equations 2 and 3 were crucial for the analytical results that followed. These are not the only reasonable choices, so bear some explanation and justification. Aside from our assumption of Bliss independence drug interaction, the other main model of null drug interactions is Loewe additivity (introduced in [59]). Loewe additivity assumes that the two drugs operate by the same mechanism of action, and therefore that the combined effect of both drugs is equivalent to the effect of either drug at their combined concentrations [19]. In this case, the mode of action and shape parameters of the two drugs must be equal, as by assumption the two drugs work interchangeably (*β*_*A*_ = *β*_*B*_ = *β, ϕ*_*A*_ = *ϕ*_*B*_ = *ϕ*). Thus, under Loewe additivity we would have that

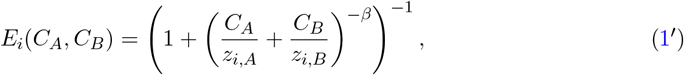

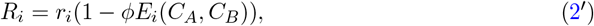

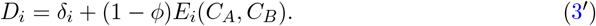

For the susceptible strain, recall that *z*_*S,A*_ = *z*_*S,B*_ = 1, and noting that *C*_*A*_ + *C*_*B*_ = *c*, we see that *E*_*S*_(*C*_*A*_, *C*_*B*_) = (1 + (*C*_*A*_ + *C*_*B*_)^−*β*^)^−1^ = (1 + *c*^−*β*^) ^−1^. That is, the effective drug concentration for the susceptible strain is only a function of the total drug concentration *c*, but not dependent on the individual drug concentrations *C*_*A*_ and *C*_*B*_. Therefore, the total number of replications 𝒩 will also be a function of *c* only, regardless of how the total drug concentration is distributed between the two drugs. Unlike with Bliss independence where intermediate drug ratios provide more efficient killing due to diminishing returns of each drug, under Loewe additivity there is no efficacy difference when using more of one drug or the other. Thus, in the Loewe additivity model, there is no tradeoff between clearing an infection faster and more mutants arising, and it is always best to use only the drug that has a lower resistance mutation rate.

In our analysis and simulations, apart from the dosing concentration and resistance mutation rate, the two drugs had identical properties. This need not be the case. If drug *A* has a higher rate at which mutations conferring resistance to it arise, but it is also more potent per unit of toxicity, it may still be preferable to use a larger dose of it than drug *B*. Moreover, the toxicity model used here is unrealistic: in reality, there is no sharp cutoff at *c* beyond which further increases in drug doses are not allowed because of catastrophic consequences and before which toxicity is zero. Instead, negative side effects are likely to be a smooth monotonically increasing function of drug concentration [60], and it could be that the two drugs have additive, antagonistic, or synergistic combined effects on total toxicity. Allowing for this greater subtlety in drug toxicity would be a valuable avenue for further research, but could complicate the mathematical analysis considerably.

One of the key weaknesses of the analytical solution presented here is that it relies on constant replication and death rates over time for all strains, whereas in reality drugs decay over time in the patient’s body. It appears that this simplification does not change the core result, however, given the introduction of pharmacokinetics in Figure S6 left the main trend unchanged. Our analytical results relied on assuming mutations confer complete resistance, where for arbitrarily large drug concentrations the mutant still achieves a positive growth rate (that is, the mutant selection window is infinitely wide). This is clearly unrealistic. Our simulation results in Figures 2 and S1 show that relaxing this assumption to allow for a realistic mutant selection window weakens but does not drastically change the result. The simulations could be extended in many ways, such as including multiple species of commensal or pathogen bacteria.

In our simulations, resource limitations led to reduced incidence of resistance mutants arising and surviving. An important effect we did not include in our analytical model, and could not detect in the simulations, is ‘competitive release’ where a strain or species that is initially limited in its population size due to competition with a fit cohabitant, can begin to grow rapidly if the competitor is eliminated [21, 61, 62]. In particular, if the susceptible bacterial population reaches a high level, then resistant mutants may struggle to grow, but once antibiotics crash the susceptible population, there is more ecological room for the resistant strains to grow. We did not observe this effect, likely because there were no pre-existing mutants in our simulations, and so even if the susceptible population crashes, there may be no resistant strain ready to fill the newly vacated niche. Thus, exploring situations with some pre-existing mutants [18] could be a valuable extension to our study.

Antimicrobial resistance is often conferred not by *de novo* mutations but through horizontal gene transfer (HGT), e.g. through the transfer of plasmids (conjugation), or the uptake of free DNA from the environment (transformation) [63, 64]. Whilst our model does not incorporate HGT of resistance genes, we believe that in some situations our results may still be applicable, at least approximately. For example, consider a scenario in which a drug-susceptible pathogen co-occurs but is not in competition with resistant commensal bacteria, and that the resistance genes can be transferred to the pathogen. In this situation, one would expect per capita rates of HGT to be roughly constant over time. (Under the commonly used mass-action assumption, the rate of HGT can be expressed as *αSD*, where *S* is the recipient and *D* the donor population size of resistant commensals, and *α* is a rate parameter.) Therefore, within our model framework, the process of HGT would be equivalent to the process of mutation (with *αD* corresponding to the mutation rate *µ*), and our results would extend to mutations acquired through HGT or a combination of both mutation and HGT. It is important to caution though that the analogy between mutation and HGT outlined above rests on the strong assumption that the number of resistant donors remains constant through time. More complex scenarios where the donor populations are also affected by the drug or interact with the pathogen population (e.g., through competition or cross-feeding) would require a new model incorporating these effects.

While these results will take time to become clinically applicable, the potential of using the (often well-characterised) resistance mutation rate in deciding on a treatment strategy is unreasonably underexplored. Even if theoretical models as abstract and (compared to reality) simple as this one cannot be directly applied in clinical settings, our results could motivate experimental efforts to corroborate them, which could in turn lead to *in vivo* tests. Our findings should in principle be straightforward to test in the laboratory. This would require assembling a set of drugs with considerably different mutation rates in some model bacteria, and challenging parallel susceptible populations with different pairs of these drugs in a variety of concentration ratios. Integrating knowledge of resistance mutation rates into pharmacological decision-making has the potential to clear more infections and minimise resistance evolution.

## Acknowledgements

We would like to thank the Engelstädter and Letten groups for discussions on this topic, in particular James Richardson and Christopher Brown. ChatGPT-4 and GitHub Copilot were used for coding assistance (but did not write any part of the manuscript).

## Data and Code Availability

All R and Mathematica code used to generate the figures and perform the symbolic manipulations, respectively, is available at https://zenodo.org/records/14197442

## Supplementary Figures

**Figure S1:**
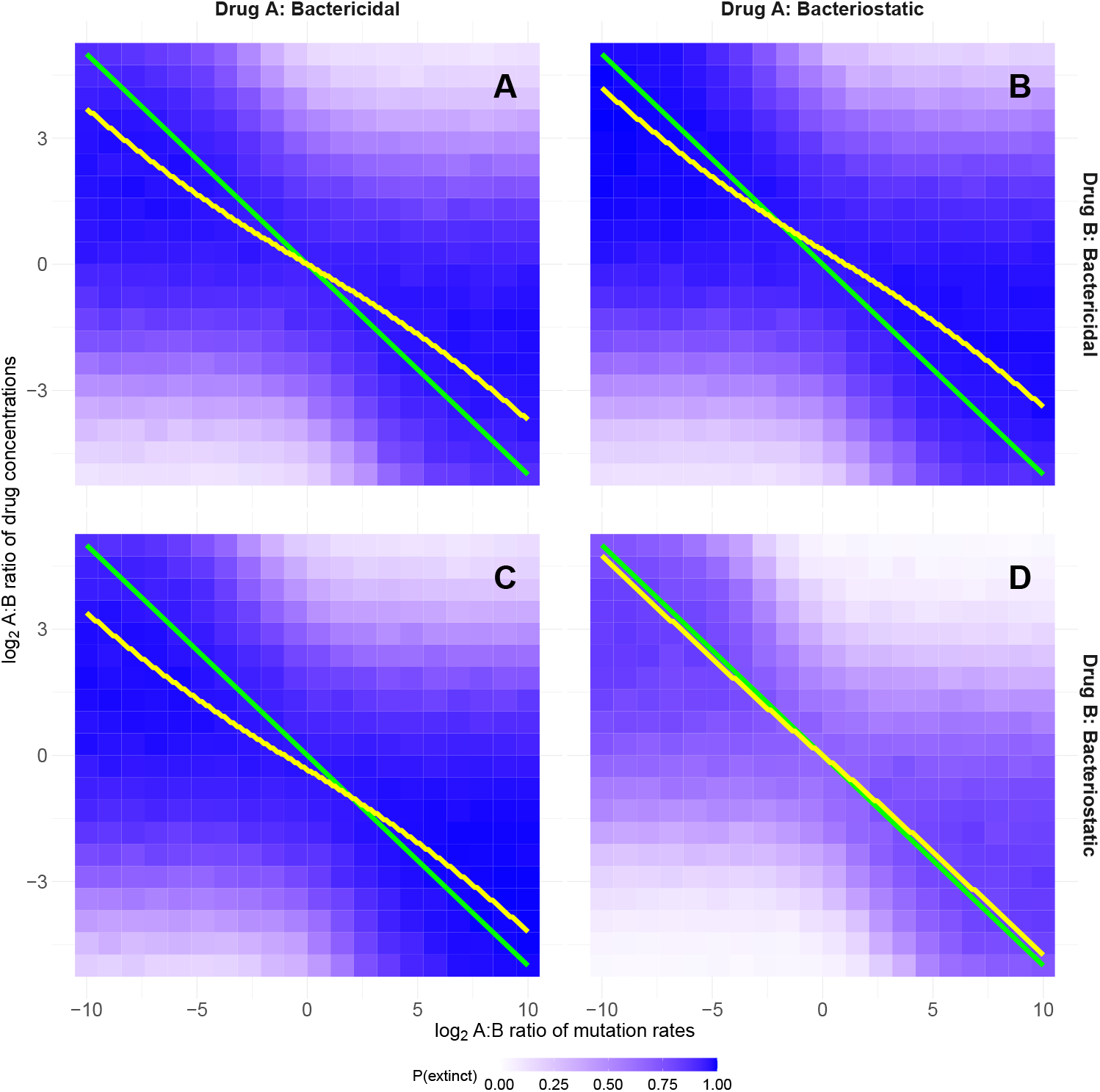
Optimal dosing with mutations conferring incomplete resistance. The parameters are identical to Figure 1 except 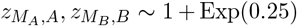, sampled independently for each run of the simulation.

**Figure S2:**
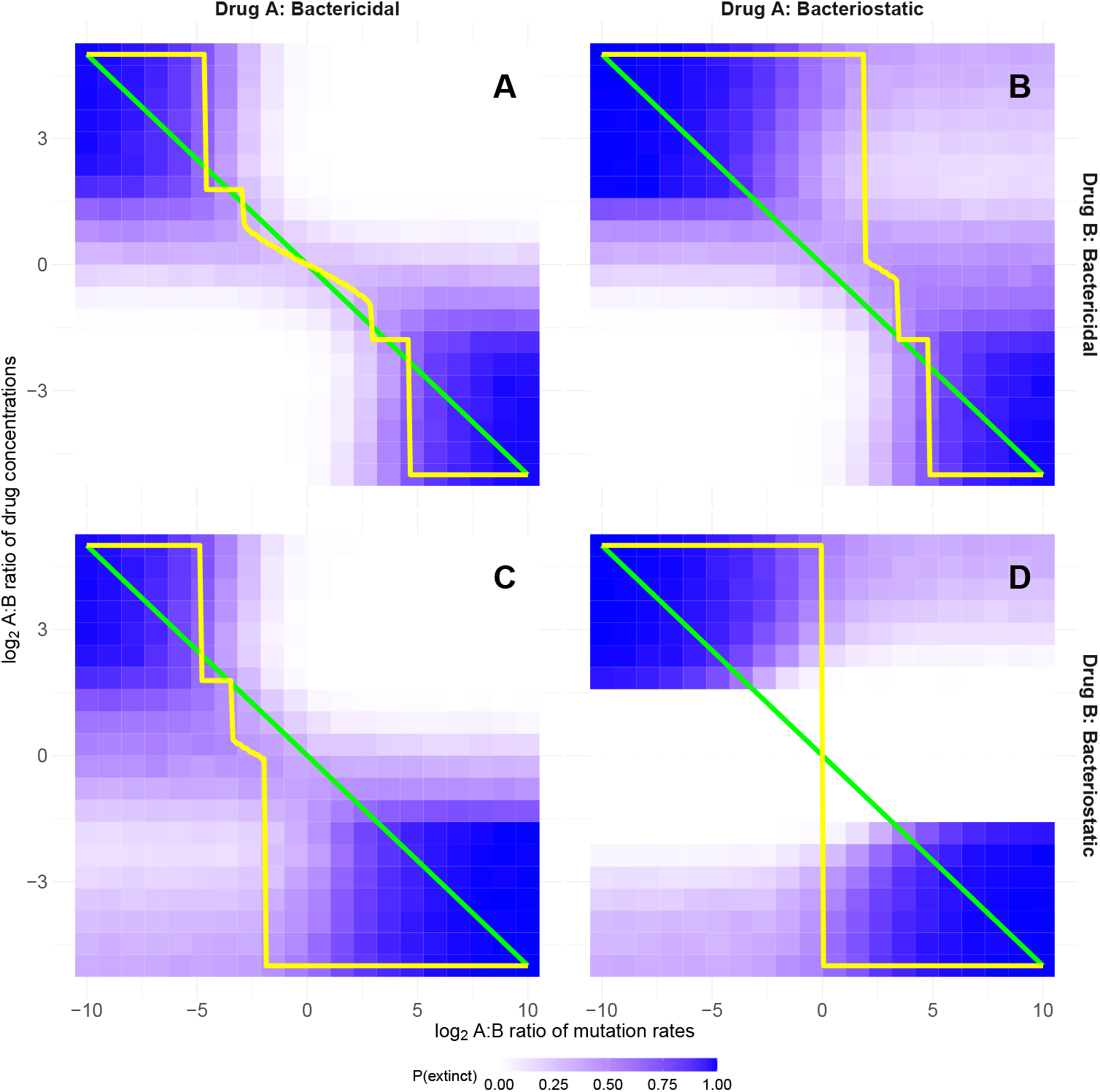
Optimal dosing with a larger shape parameter sometimes entails using solely one drug. The parameters are identical to Figure 1 except *β*_*A*_ = *β*_*B*_ = 3 and *δ* = 0.19.

**Figure S3:**
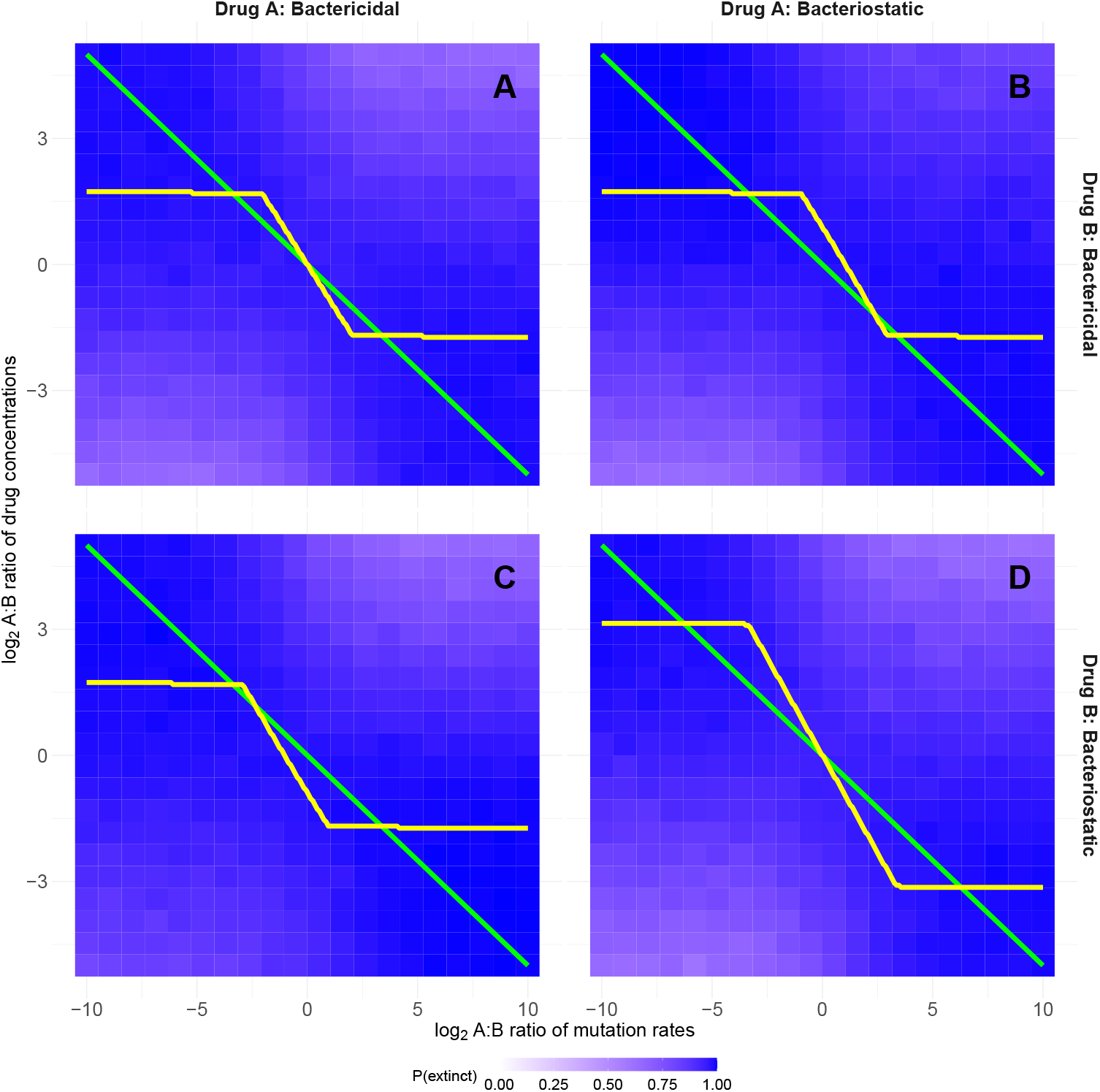
Optimal dosing with a smaller shape parameter. The parameters are identical to Figure 1 except *β*_*A*_ = *β*_*B*_ = 0.2, *δ* = 0.47. In the top-left and bottom-right corners of each panel, *P*_*E*_ = 1 and so the precise drug concentration ratio used does not matter, as long as it is beyond some threshold. This is reflected in the z-shaped truncated yellow lines that do not require arbitrarily skewed drug dosing ratios.

**Figure S4:**
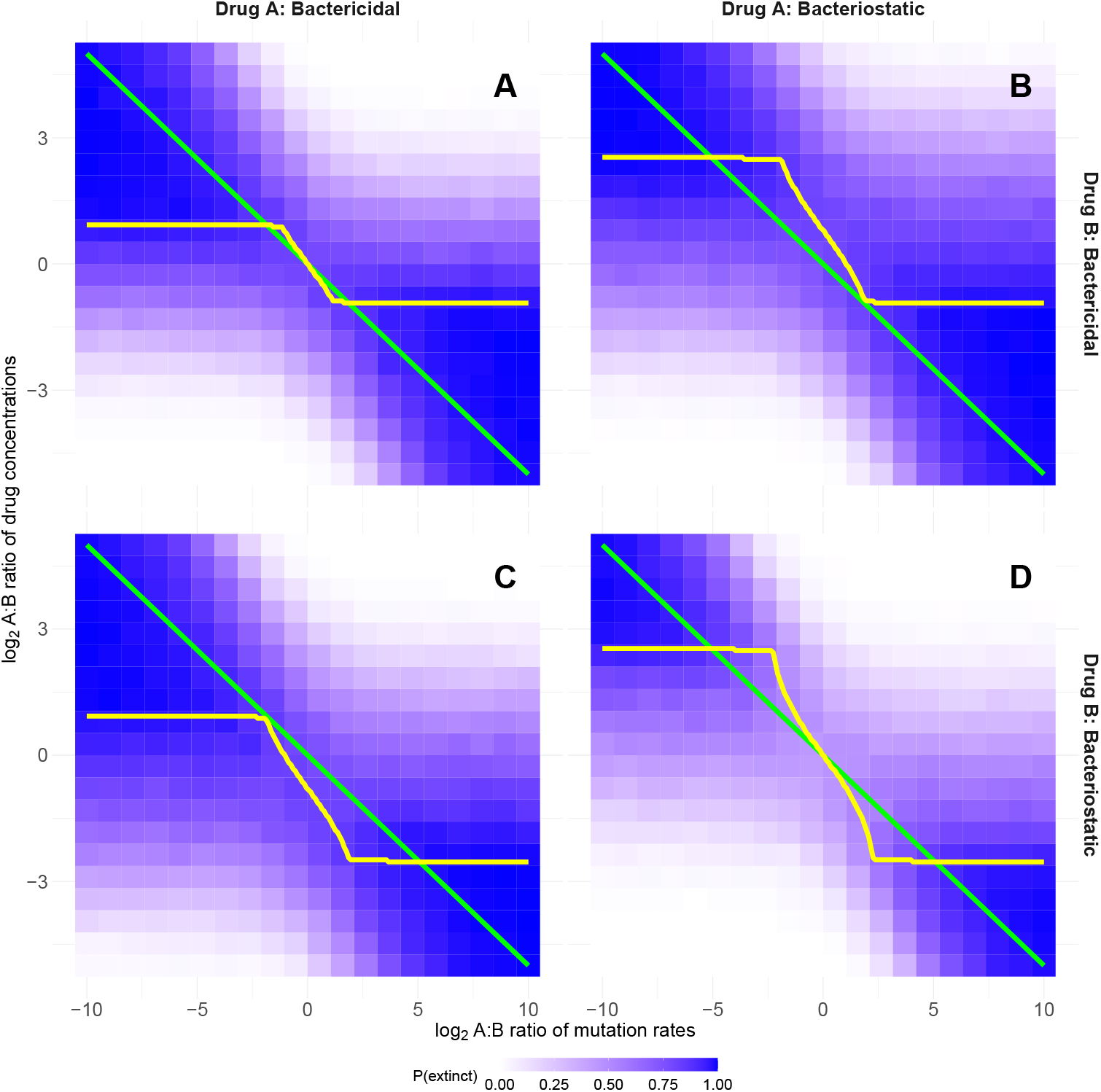
Optimal dosing with costs of resistance. The parameters are identical to Figure 1 except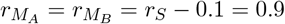.

**Figure S5:**
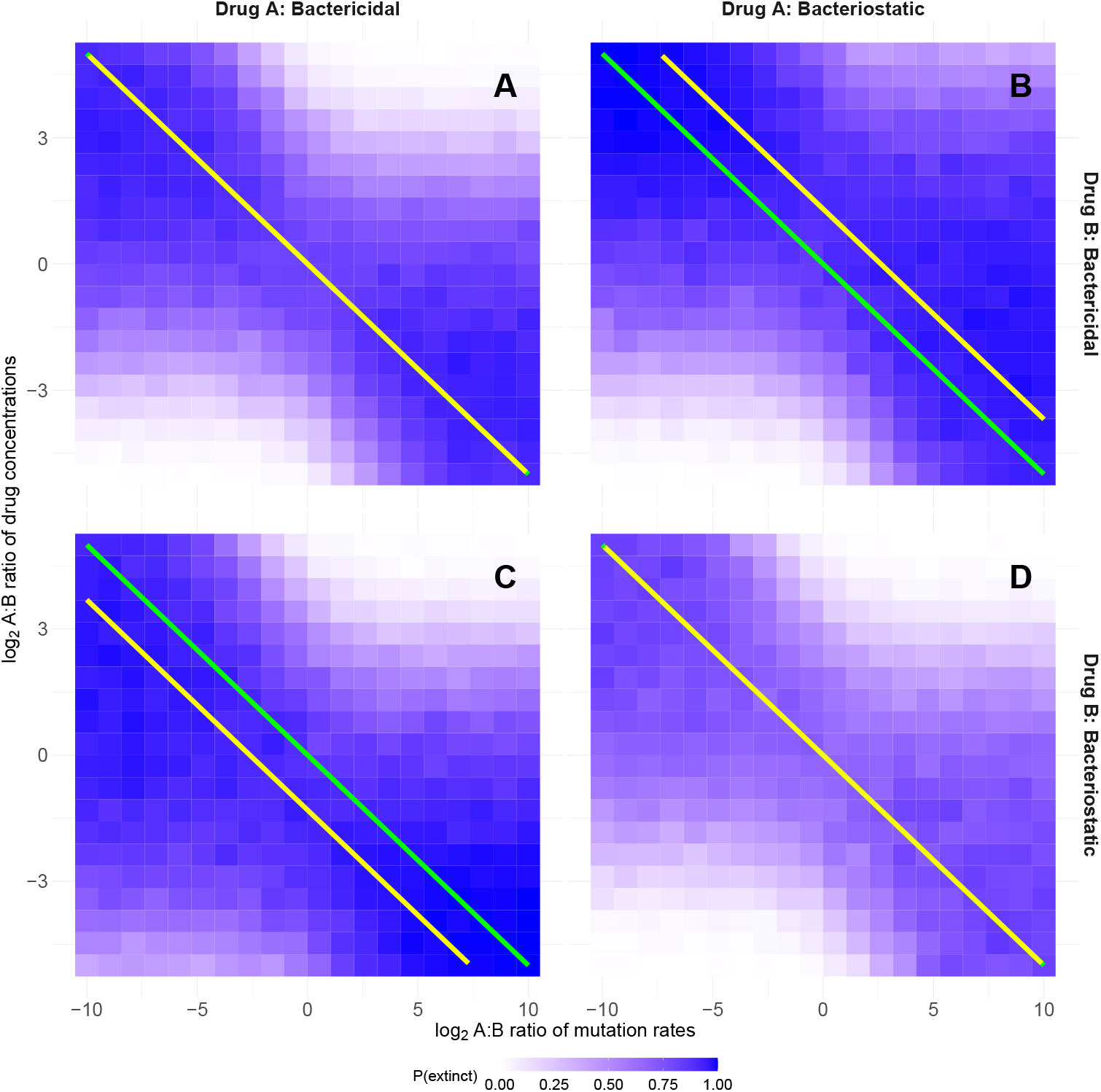
Optimal dosing with higher drug concentrations. The parameters are identical to Figure 1 except 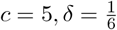.

**Figure S6:**
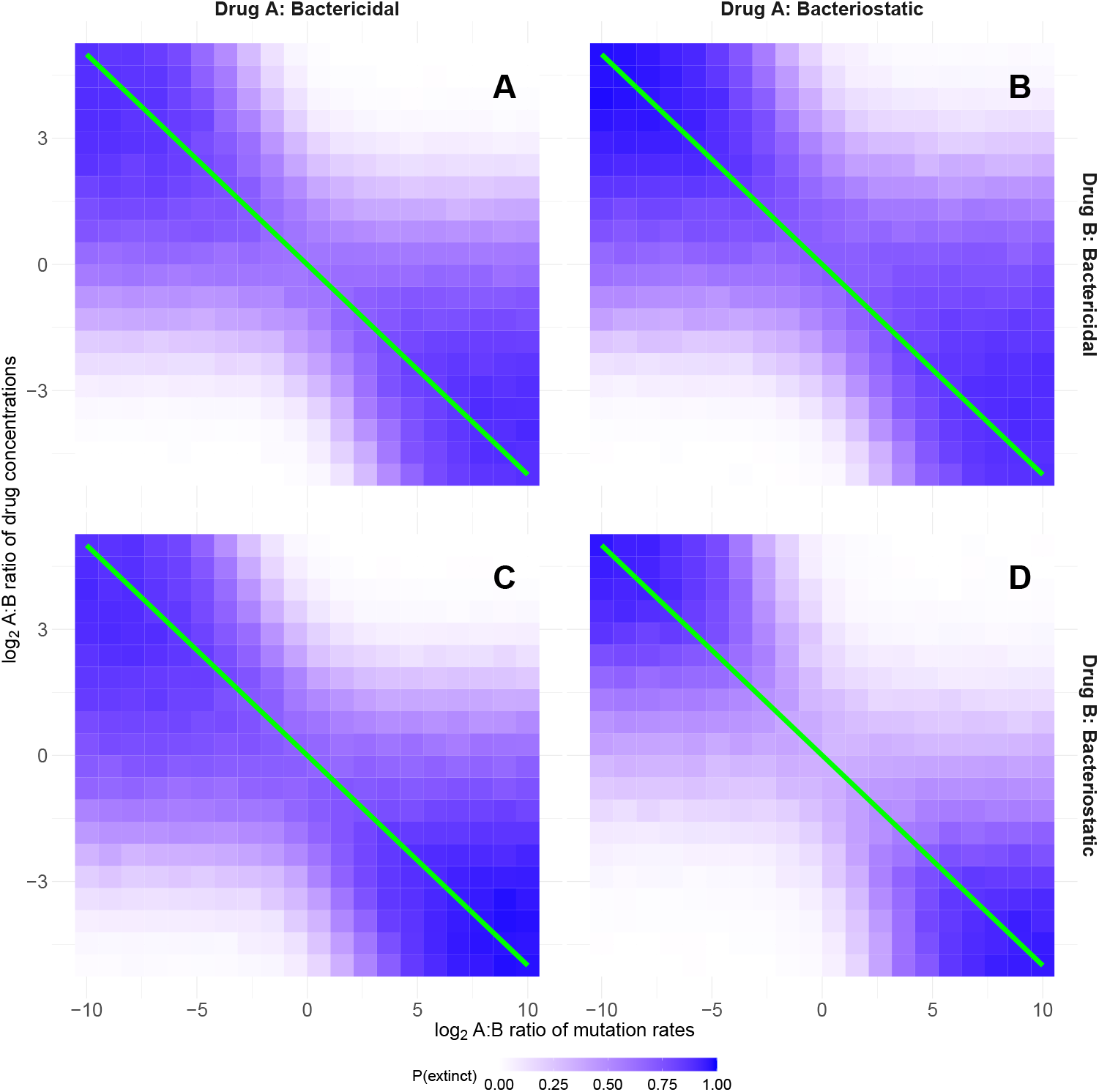
Optimal dosing with pharmacokinetics. The parameters are identical to Figure 1 except a drug decay rate of 0.15 has been introduced with doses every 12 hours of both drugs, meaning that *e*^−0.15×12^ = 17% of the previous dose remains at the next dose. To compensate for the drug decaying, the intrinsic death rate has been increased by 0.2 to *δ* = 0.53. The yellow theory lines are not shown here, as the theoretical analysis only dealt with constant drug concentrations. The original green lines are still shown for comparison.

**Figure S7:**
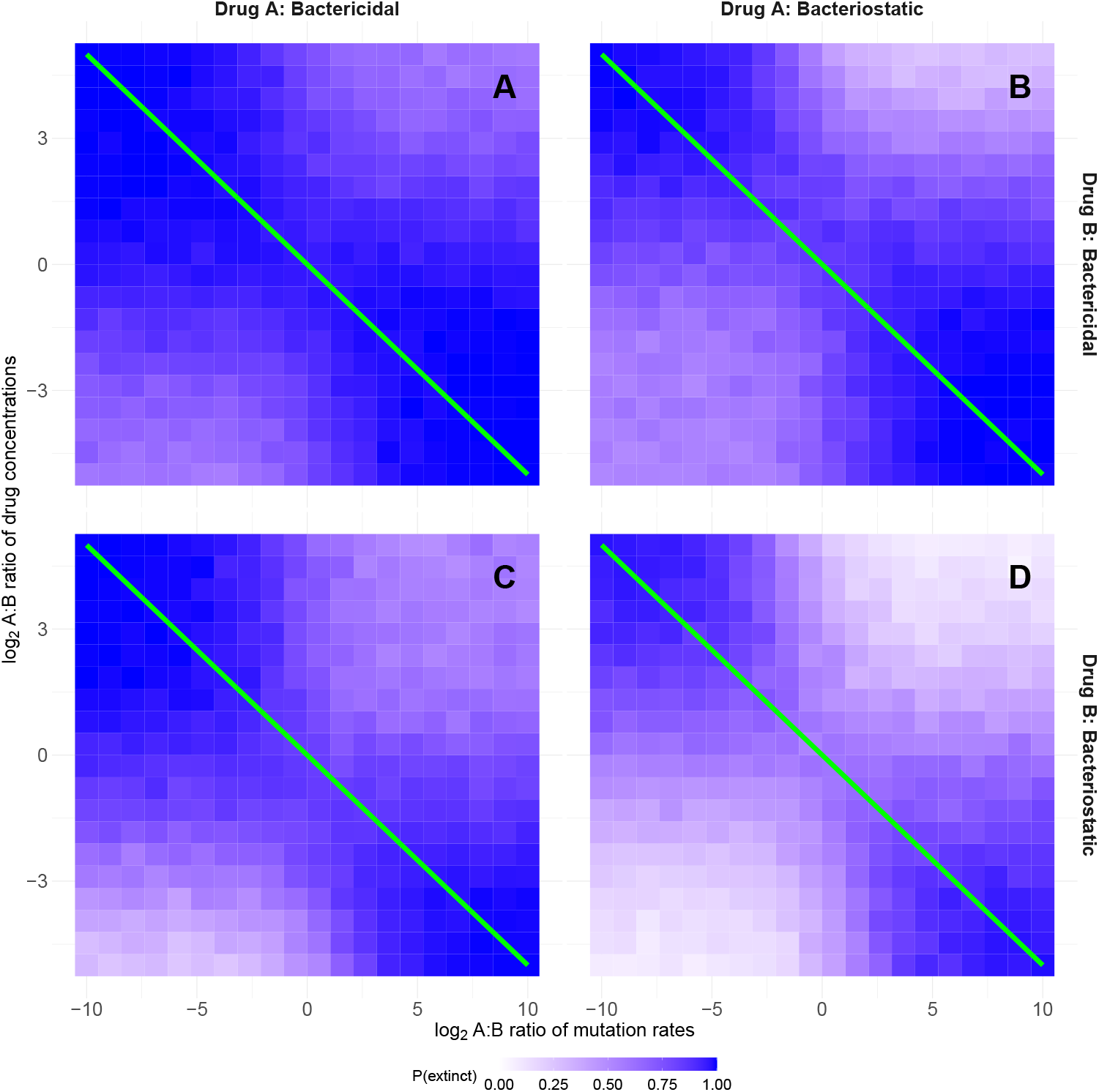
Optimal dosing with resource constraints. The parameters are identical to Figure 1 except growth is now modelled as being limited by a single rate-limiting resource, with an initial concentration of 10^9^ units, where one unit is consumed per bacterial replication, and a constant resource influx of 10^8^ h^−1^. The yellow theory lines are not shown here, as the theoretical analysis assumed unconstrained growth. The original green lines are still shown for comparison.

